# Plakins are involved in the maintenance of epithelial polarity

**DOI:** 10.1101/2023.10.16.562500

**Authors:** Juliana Geay, Yoran Margaron, David Gentien, Fabien Reyal, Alain Puisieux, Laurent Blanchoin, Laurent Guyon, Manuel Théry

## Abstract

As epithelia are the interface between the organism and the external environment, they are often subject to damage and must be frequently renewed. However, maintaining epithelial integrity during this renewal is challenging, and loss of cell polarity is a potent inducer of tumorigenesis. In this study, we used transcriptomic data from breast cancer cells at different stages of tumor development to identify molecular changes associated with the early stages of tumor transformation. We correlated these protein expression profiles with either cell polarity defects or cell progression along the epithelial-to-mesenchymal transition (EMT). We identified plakins, namely epiplakin (EPPK1), desmoplakin (DSP) and periplakin (PPL), that were downregulated in cells that had lost their epithelial polarity and also downregulated in cells that had progressed through EMT. We further tested them experimentally by knocking down their expression in a non-tumorigenic epithelial breast cell line (MCF10A). We demonstrated their causal role in the loss of polarity, as revealed by the misorientation of the nuclear centrosome vector. We also found that vimentin, a marker of EMT, was overexpressed in plakin knocked-down cells, suggesting that plakins may have both a structural and a regulatory role in maintaining the epithelial state.

## Introduction

Epithelia lines the interface between organs and external environment. Their morphogenesis involves tightly regulated coordination of multiple cytoskeleton components (Schöck and Perrimon, 2002; Roignot et al., 2013). They are intensively renewed as they are exposed to numerous physical and chemical stress (Darwich et al., 2014; Okumura and Takeda, 2017). It is challenging to renew such structures in which tissue cohesion is key for their function. Defective maintenance of tissue integrity can lead to epithelial cancers called carcinoma (Hinck et al., 2014).

The loss of epithelial cell polarity is a major cause of cancers (Wodarz and Näthke, 2007). Cell polarity cues appeared to act as oncogenes (Lee and Vasioukhin, 2008). Whether the loss of polarity is a cause or consequence of tumoral development has been a long-standing question in the field (Muthuswamy and Xue, 2012). Tumoral progression is a complex process that associates uncontrolled cell growth and tissue disorganization. To better understand its origin it is key to figure out what are the earliest signs of tumor development.

The dissemination of breast carcinoma, as for numerous cancers, proceeds through an epithelial-to-mesenchymal transition (EMT) (Brabletz et al., 2018; Saitoh, 2018; Tsai and Yang, 2013). Loss of epithelial polarity is an early sign of EMT (Jung et al., 2019). In particular, centrosome disconnection from junctions and repositioning at the cell center precedes junction disassembly (Burute et al., 2017). So arguably centrosome mispositioning can be considered as one of the earliest signs of tumoral development. It is unclear whether centrosome mispositioning is a consequence of earlier deleterious changes in cell organization or a cause of further downstream disorganizations. Both are likely true, since the process regulating centrosome position integrates the contributions of several cellular structures (focal adhesions and intercellular junctions, actin network, nucleus and endomembrane networks) (Haupt and Minc, 2018; Jimenez et al., 2021), and, in return, the position of the centrosome will affect the spatial distribution of microtubules and thereby the transport of proteins and signals throughout the cytoplasm (Barlan and Gelfand, 2017). It is therefore key to understand the molecular and cellular mechanisms leading to centrosome mispositioning in early stages of tumoral transformation.

Numerous transcriptomic data sets have been obtained from cells lines at various stages of tumoral transformation and constitute a rich source of information. However, they provide measurements about the cellular content, in terms of RNAs, but nothing about the organization of cellular structures. How could we relate these molecular measurements to the architecture and polarity of cells?

The maintenance of epithelia and cell architecture under constant renewal is a complex morphogenetic process, which is multifactorial and dynamic (Roignot et al., 2013). It is therefore difficult to manipulate it experimentally and obtain a reproducible multicellular conformation that is amenable to precise quantification in order to detect subtle changes in early stages of tissue transformation. To that end we developed a minimal polarized system made of a pair of epithelial cells, geometrically controlled in a non-moving steady state on a micropattern of fibronectin (Tseng et al., 2012). This minimal system recapitulates the epithelial segregation of cell-cell and cell-ECM adhesion (Burute et al., 2012) and therefore direct centrosome positioning and epithelial cell polarity (Burute et al., 2017).

In this study, we analyzed two sets of transcriptomic data from breast cancer cell lines in order to identify potential regulators of centrosome positioning and epithelial polarity. We then tested three candidates experimentally on doublets of non-transformed human mammary epithelial cells.

## Results

### Variation of gene expression with the loss of epithelial cell polarity

Triple negative breast cancer cells (TNBCs) are resistant to hormone treatment and thus associated with poor prognosis (Lehmann et al., 2011). To help the identification of new treatments, the transcriptomics profiles of TNBCs were measured and made accessible to the community (Rody et al., 2011). Importantly, a key feature of TNBCs malignancy is that they have initiated EMT and thus are prone to induce the formation of metastases (Hudis and Gianni, 2011; Dent et al., 2009; Sarrió et al., 2008; Jang et al., 2015). The invasion step being initiated by the dissociation and escape from the primary tumor, we hypothesized that the epithelial polarity of TNBCs was impaired. Furthermore, we thought that a quantitative description of this structural defect in various lines of TNBC would allow us to relate the degree of polarity loss to specific variations of gene transcription.

We thus first measured the epithelial polarity of 11 TNBC lines by cultivating cell doublets on H-shaped fibronectin-coated micropatterns over 24 hours (Figure 1A). The geometry of the micropattern imposed the shape and architecture of cells, and notably the reproducible position of the intercellular junction along the vertical axis bisecting the H (Tseng et al., 2012). The position of the centrosome with respect to the nucleus and the inter-cellular junction can be taken as a proxy for the orientation of cell polarity (Burute et al., 2017) (Figure 1B). To capture the behavior of the entire population in a single metric, we measured the proportion of cells in which the centrosome was clearly off-centered toward the inter-cellular junction and named this value the “polarity score” (Figure 1B). We measured it in three independent experiments for the 11 TNBC cell lines and the non-tumorigenic epithelial breast cell line MCF10A as a control. The polarity scores of all TNBCs were remarkably lower than in MCF10A (Figure 1C). However, these low scores were not identical for all TNBCs and variations could be related to the specific transcriptomic profile of each cell line.

**Figure 1.**
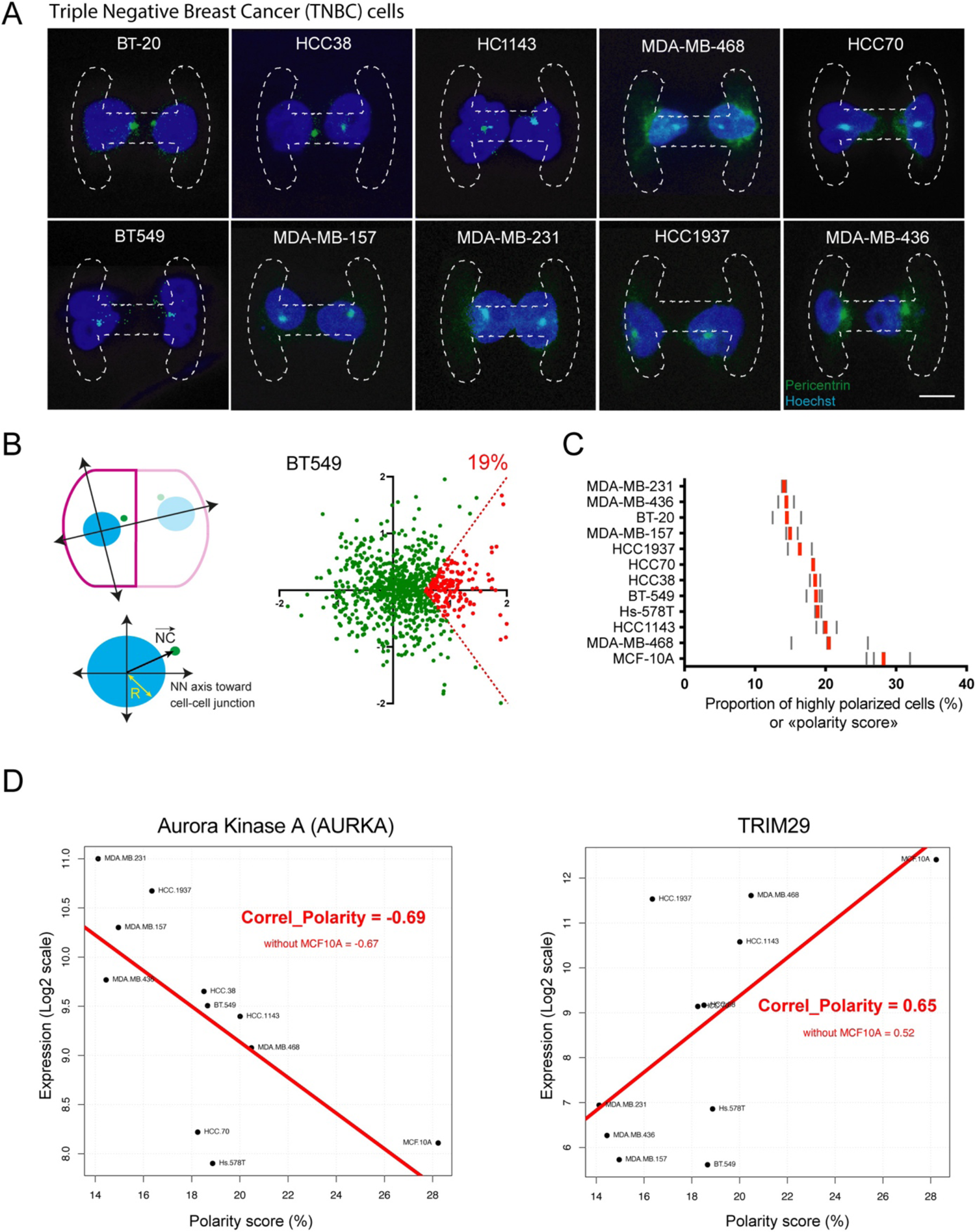
Variation of gene expression with the polarity of TNBC cells. A - Polarity of TNBC cells. Representative images, from maximum projections of Z-stacks, showing cell doublets cultured during 24 hours on H-shaped micropatterns (drawn with dashed lines). Cells were stained with pericentrin (green) and Hoechst (blue) for centrosome and nucleus location, respectively. Scale bar represents 20 µm. B - The scheme shows the centrosome coordinates with respect to the center of mass of the nucleus: x-axis corresponds to the nucleus-nucleus (NN) axis passing through the center of two nuclei. The distance from nucleus center to centrosome is normalized by the nucleus radius. The graph shows the spatial distribution of centrosome position with respect to the nucleus in the reference axis represented in the scheme in the case of the BT549 cells. The red sectors correspond to cells that are further considered as “highly polarized”. C - The histogram represents the “Polarity score”, ie the average proportion of “highly polarized cells” in each TNBC cell lines over three independent experiments. Each grey bar represents the proportion of highly polarized cells measured over more than 200 cells per experiment. The red bar represents the averaged polarity score over three independent experiments. D - Examples of two genes whose expression level is highly correlated to the “Polarity score” of the 11 TNBC cell lines and the MCF10A. Aurora Kinase A is an example of a negative correlation, TRIM29 is an example of a positive correlation. “Correl_Polarity” indicated the value of the correlation between protein expression levels and the Polarity score. MCF10A could bias this correlation since they have a high polarity score, so we also measured the correlation coefficient without taking this cell line into account.

For each gene, we measured the correlation coefficient between the amount of transcripts in each cell line and their polarity score. Interestingly, genes for which expression was negatively correlated with the polarity scores were enriched in cell division genes, whereas the positively correlated ones were enriched in intercellular adhesion and differentiation genes (listed in Table S1). As expected from literature we identified some genes whose expression level was negatively correlated to the polarity scores, ie they were less expressed in the most polarized cells, such as Aurora Kinase A (Wirtz-Peitz et al., 2008), and others which were positively correlated, ie they were more expressed in most polarized cells, such as TRIM29 (Liu et al., 2012) (Figure 1D). For those “hits” we tested whether the value of the correlation was not influenced by their expression in MCF10A which is a bit of an outlier in the group.

### Variation of gene expression as cells progress along EMT

We then searched for a second and independent way to reveal the genes involved in the maintenance of epithelial polarity. The apico-basal polarity of epithelial cells is progressively lost as cells progress along EMT (Moreno-Bueno et al., 2008). Cells progress in this dedifferentiation and redifferentiation pathway depending on their degree of tumoral transformation (Su et al., 2020). This complex landscape was clarified by a broad investigation of the variations of gene expression in a non-transformed human mammary epithelial cells (HME) in response to the over or down expression of key EMT transcription factors (Twist 1/2, Zeb1/2, TGF-β) and cancer genes (p53, Ras) (Morel et al., 2012). To relate those changes to cell progress along EMT, we calculated for each modified cell line an “EMT score” based on a molecular signature made of 150 down-regulated genes and 89 up-regulated genes that characterize the core changes common to various forms of EMT (Taube et al., 2010) (see Materials and Methods). HME cell lines were ranked based on this EMT score (Figure 2A). Non-surprisingly, the MDA-MB-157 cell line which served as a control of highly transformed breast cancer cell line, scored high in the rank, as well as modified cell lines expressing EMT transcription factors in combination with Ras. As we did previously for TNBCs transcriptomes and polarity scores, we measured the correlation coefficient between the amounts of transcripts in each cell line and their EMT score (listed in Table S2). Genes for which expression is anticorrelated with EMT score are enriched in associated proteins localized in various organelles connected by the cytoskeleton (cornified envelope, cell junctions, desmosomes, mitochondrion, etc.) and participate in cadherin-binding and keratinocyte differentiation. Genes with positive correlation are enriched in associated proteins localized in focal adhesion, cytoskeleton, and endoplasmic reticulum, and participate in cell adhesion (but not cell-cell adhesion) and migration. As expected from literature, we found typical genes that were negatively or positively correlated to the EMT scores, such as Keratin15 (Zhong et al., 2021) or collagen XV (Yao et al., 2022), suggesting that the regulation of their expression level was part of the intra-cellular reorganization that accompanies EMT.

**Figure 2.**
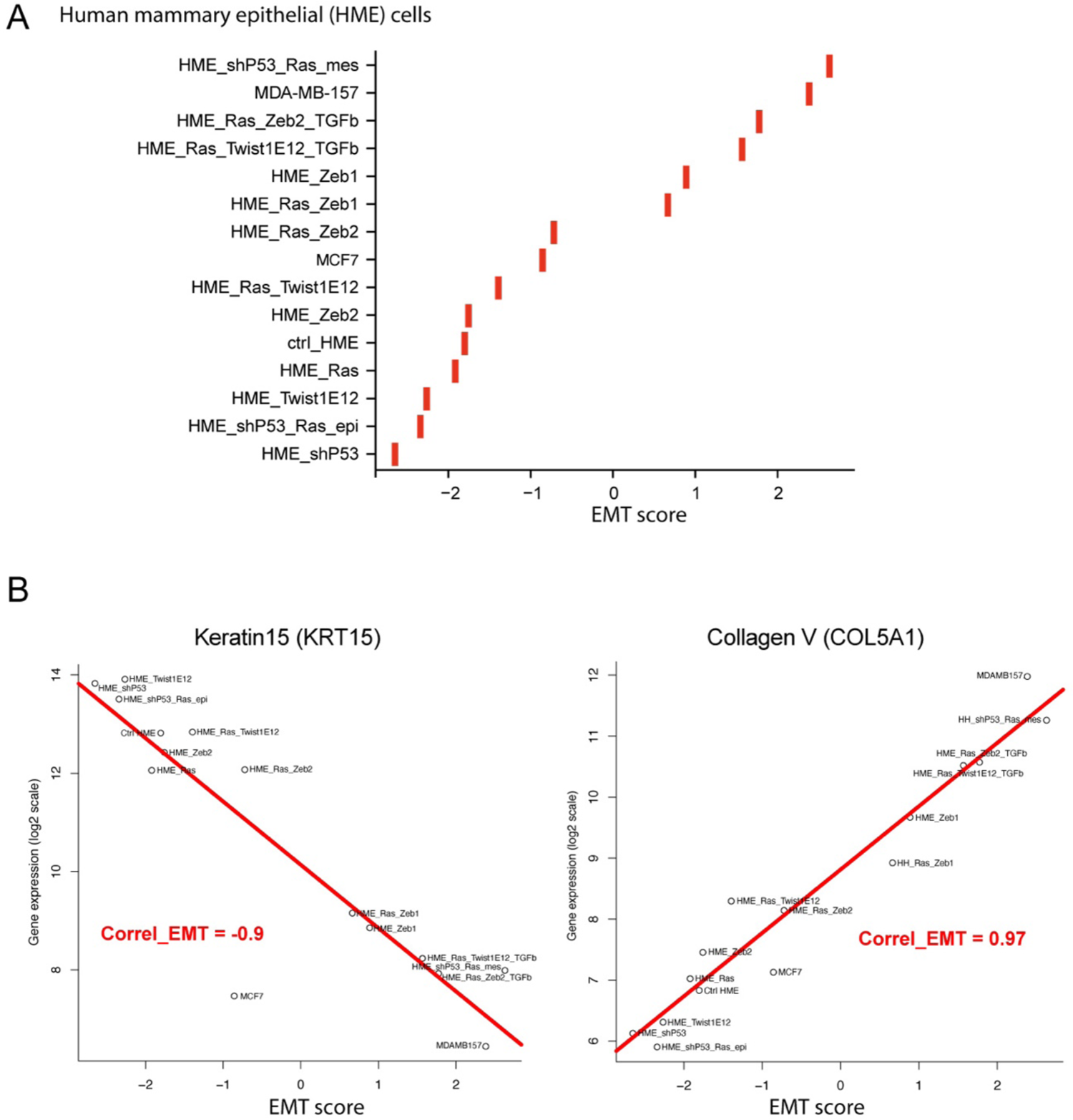
Variation of gene expression as HME cells progress along EMT. A - Calculation of the “EMT score” of HME cell lines over-expressing various transcription factors or oncogenes. The EMT score has been designed based on the EMT signature described in (Taube et al., PNAS, 2010) (see Methods). B - Examples of two genes whose expression level is highly correlated to the “EMT score” of the HME cell lines. Keratin 15 is an example of a negative correlation, Collagen V is an example of a positive correlation. “Correl_EMT” indicated the value of the correlation between protein expression level and the EMT scores.

### Plakins at the intersection of EMT, polarity and microtubules

In these two lists of selected hits, we then searched for potential candidates that would affect the internal organization of the microtubule network architecture, and thereby affect centrosome position and epithelial cell polarity. We selected a hundred of proteins, including microtubule stabilizing and destabilizing proteins (CLIP, CLASP, CAMSAP, MAP2, Tau, stathmin, katanin, spastin…), all kinesins and dynein, as well as cytolinkers such as plakins and spectrins (Table S3). We then plotted the values of their correlation with EMT against their correlation to polarity that we measured previously. The graph showed an overall negative trend, which means that many genes that were positively correlated with EMT in HME were instead negatively correlated with epithelial polarity of TNBCs (Figure 3). This was expected and confirmed that the progression along EMT is associated with a loss of epithelial polarity (Moreno-Bueno et al., 2008). Several key regulators of epithelial polarity such as CAMSAP3 (Noordstra et al., 2016; Toya et al., 2016) displayed highly positive correlation with polarity and highly negative correlation with EMT. On the opposite trend, MAP1B appeared anti - correlated to epithelial polarity, consistent with its role of microtubule destabilizer and regulator of neuronal cell polarity (Tortosa et al., 2013). Kinesins were distributed all over the graph but, as previously shown, the expressions of KIFC3 and KLC3 showed high and opposite correlation with epithelial polarity: KIFC3 being negatively correlated (Lu et al., 2023) whereas KLC3 was positively correlated to it (Fustaino et al., 2017). Interestingly, three plakins, namely epiplakin (EPPK1), desmoplakin (DSP) and periplakin (PPL) all appeared as clear outliers (Figure 3). The expression of each of these plakins, together with envoplakin (EVPL), were negatively correlated to the EMT score of HME and positively correlated to the Polarity score of TNBC (Figure S1). Although a role in epithelial polarity maintenance was not completely surprising for these gigantic proteins that connects intercellular junctions to microtubules and intermediate filaments (Bouameur et al., 2014; Suozzi et al., 2012), such a strong coupling of several members of the plakin family with both EMT and polarity was not expected.

**Figure 3.**
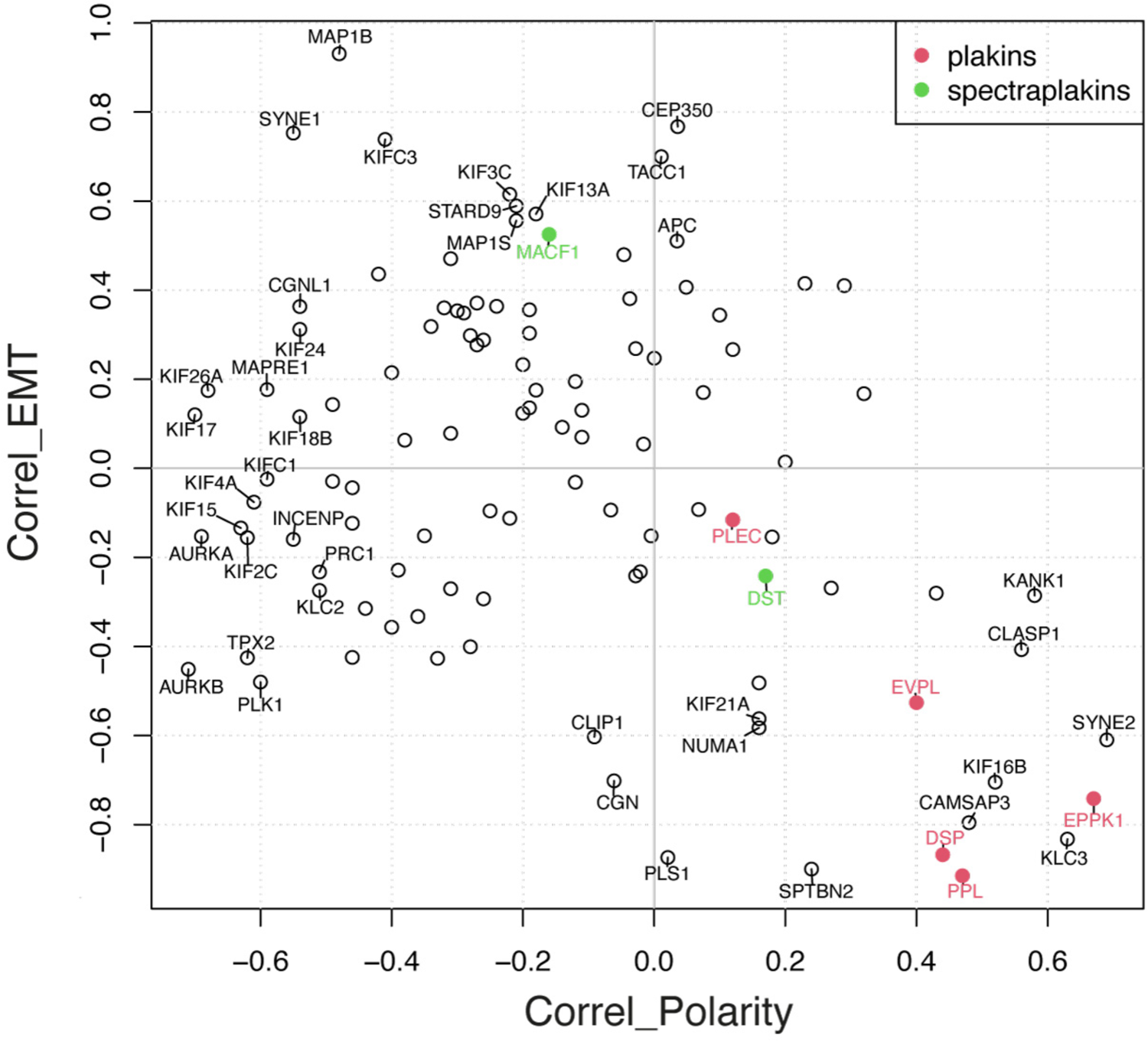
Plakins at the intersection of EMT, polarity and microtubules. This graph represents the correlations values of the levels of expression of the protein with respect to the EMT score of HME cells (see figure 2) and with respect of the Polarity score of TNBC cells (see figure 1) for a hundred of microtubule associated proteins. Plakins are shown in red.

### Knockdown of plakins induce a loss of epithelial polarity

To directly test the role of plakins in the regulation of epithelial polarity, we knocked - down the expression of epiplakin, desmoplakin and periplakin individually. We tested several siRNA sequences and retained two for each that clearly downregulated the expression of the targeted proteins as attested by the immunostaining of western blots (Figure 4A and Material and Methods). We also found a marked disappearance of their cytoplasmic localization (Figure 4B). Interestingly, knocked-down cells in culture displayed abnormal shapes, being more elongated and less cohesive (Figure 4C).

**Figure 4.**
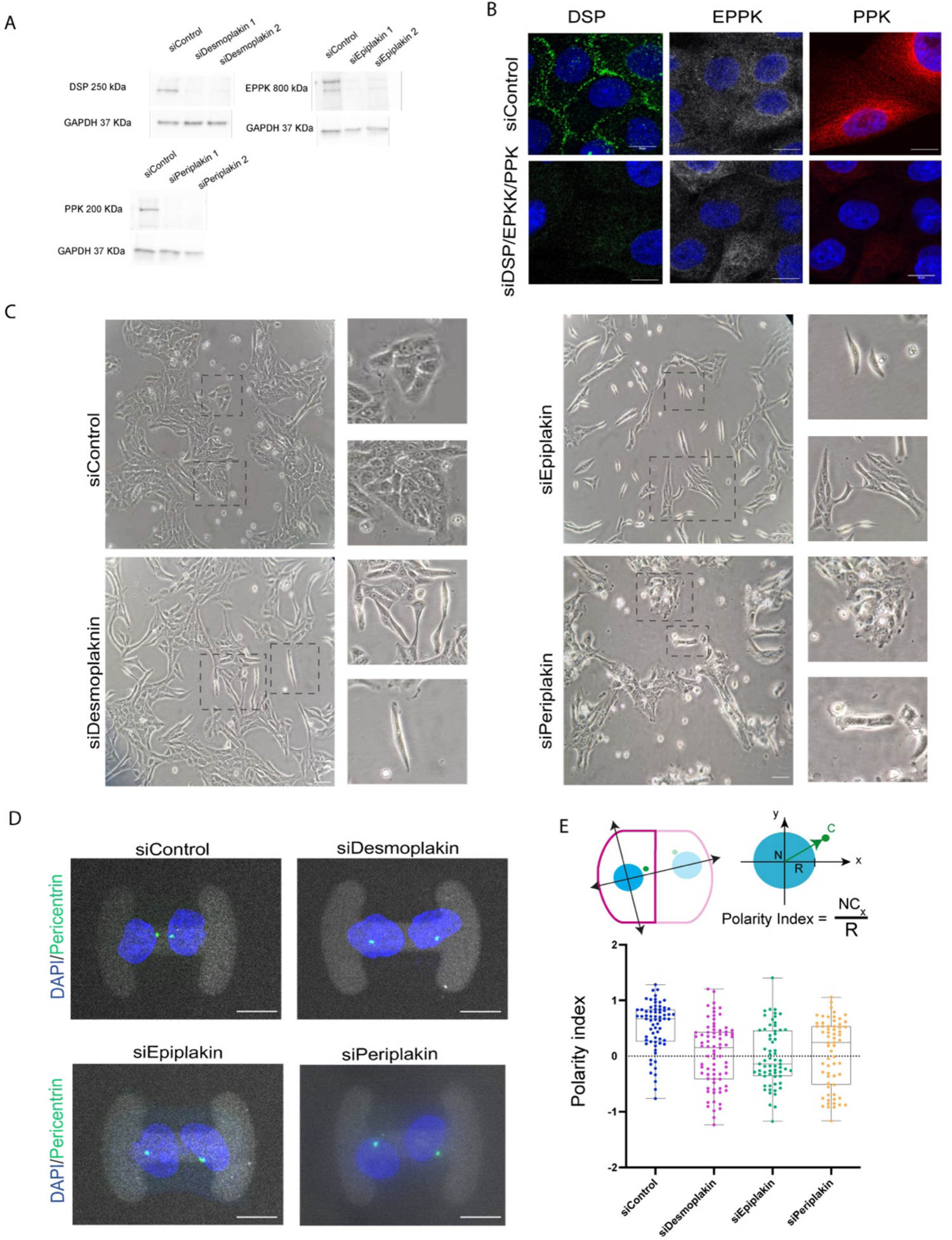
Knockdown of plakins induce a loss of epithelial polarity A - Western-blots of MCF10A cells treated with a control siRNA and siRNA against Desmoplakin (DSP), epiplakin (EPPK) and periplakin (PPK). B - Confocal images of cells stained with Desmoplakin, Epiplakin and Periplakin in MCF10A cells treated with siDSP, siEPPK, siPPK. Images correspond to a maximum intensity projection of 6 slices, spaced by 1 micron. Bar, 10 μm. C - Brightfield images of MCF10A cells treated with siControl, siDSP, siEPPK, siPPK showing morphological differences in the depleted cells and single cell emergence. Images were acquired by bright-field microscopy using a 10× objective. Scale bar correspond to 10 μm. D - Confocal images of MCF10A cells treated with siControl, siDSP, siEPPK, siPPK plated on H micropatterns and immunostained with antibodies against pericentrin to label the centrosome and stained with DAPI to label the nucleus. The fluorescent fibrinogen of the micropattern is shown in grey. Images correspond to a maximum intensity projection of 6 slices, spaced by 1 micron. Scale bar correspond to 10 μm. E - The scheme describes the detection of centrosome position and calculation of the corresponding polarity index. Graph represents polarity index measurements of centrosome positioning in MCF10A cells treated with siControl, siDSP, siEPPK, siPPK. When 0.5<PI<2, centrosome is off-centered toward the cell-cell junction; when −2<PI< − 0.5, centrosome is off-centered toward the ECM. Horizontal lines indicate the median value for each cell type. Represented data shown are from three independent experiments, for which n was between 30 and 40 cells for each cell type. ****p < 0.0001 by the Mann-Whitney U-Test.

To characterize cell ability to polarize and orient the organization of their microtubule network, independently of these cell shape changes, we again used the micropatterning of cell doublets on H shapes. We measured the cell polarity index (Burute et al., 2017), ie the coordinate of the centrosome along the nucleus-junction axis for control MCF10A and knocked-down cells (Figure 4D and 4E). As expected for properly polarized cells, MCF10A displayed positive polarity index, with an average value of 0.5, similar to previous measurement in our previous studies (Burute et al., 2017). By contrast, the polarity index of cells knock-down for epiplakin, desmoplakin or periplakin were close to zero, meaning that their centrosomes were randomly oriented around the nucleus, and independent of the position of the intercellular junction. Similar measurements were obtained by treating cells with the second siRNA sequence for each plakin (Figure S2). These results confirmed that epiplakin, desmoplakin and periplakin are all potent regulators of epithelial cell polarity.

### Impact of plakins on cytoskeleton networks

We finally measured whether cytoskeleton networks were impacted by the loss of plakins. We first analyzed the actin network and found that the loss of periplakin impaired significantly the ability of cells to form inter-cellular junctions, as attested by the shortening of these junctions (Figure 5A). However, we found no effect of the loss of desmoplakin or epiplakin on junction length (Figure 5A), which was consistent with previous reports showing that E-cadherin expression was not altered in mouse knocked-out cells, although these became invasive (Chun and Hanahan, 2010). This suggested that centrosome mispositioning in these cells was not due to weaker junctions but rather a downstream consequence of a defective connection between junctions and the microtubule network.

**Figure 5.**
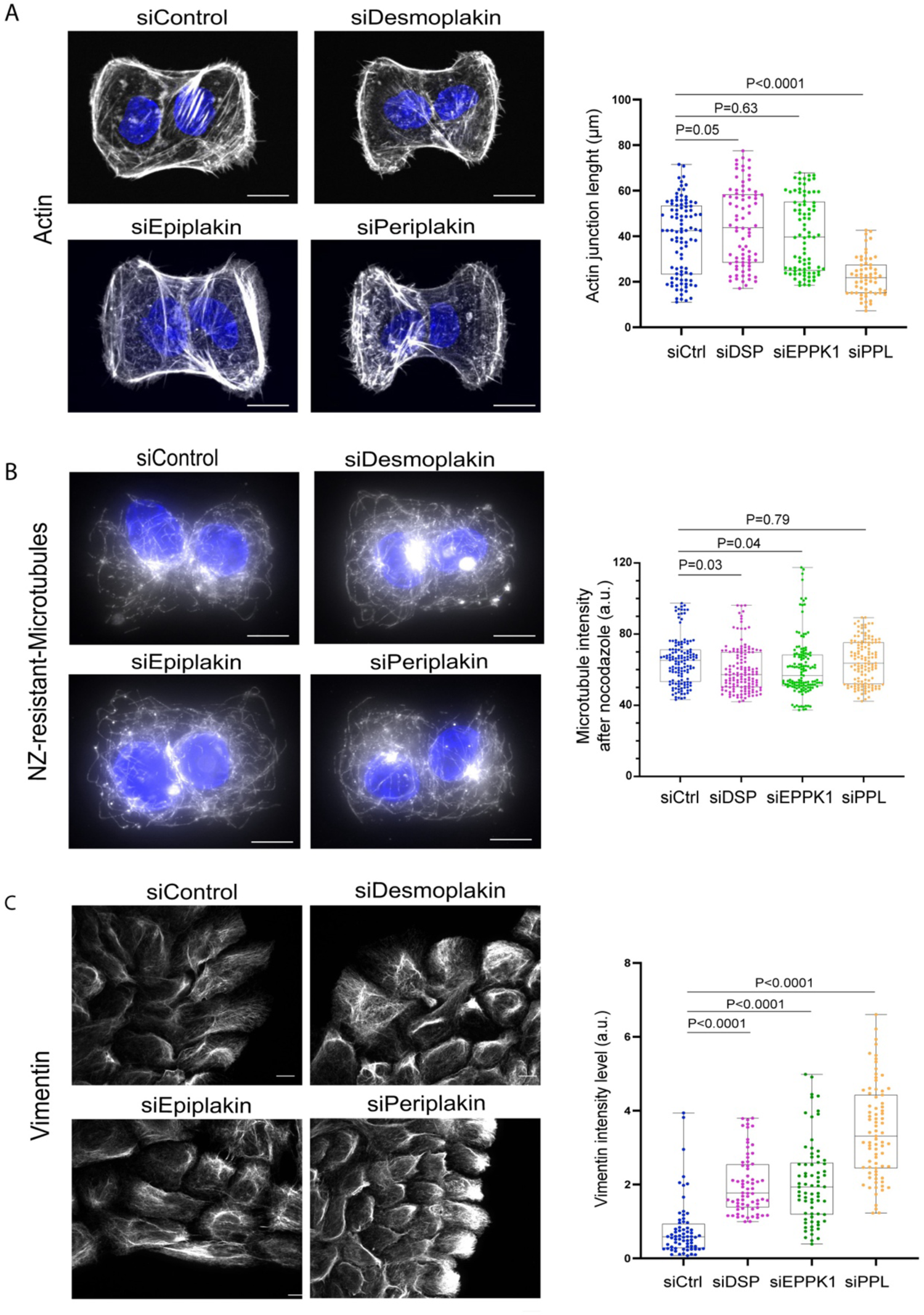
Impact of plakins on cytoskeleton networks. A - Confocal images of MCF10A cells treated with siControl, siDSP, siEPPK, siPPK and stained with phalloidin to reveal the architecture of the actin network. Images correspond to a maximum intensity projection of 6 slices, spaced by 1 micron. Scale bar correspond to 10 μm. Graph represents the length of the intercellular junction. Measurements were obtained from three independent experiments, for which n was between 10 and 20 cells for each condition. Indicated P-values were obtained after performing Mann-Whitney U-Tests. B - Confocal images of MCF10A cells treated with siControl, siDSP, siEPPK, siPPK and further treated with 10µM Nocodazole for 5 minutes to reveal only the most resistant microtubules. Fixed cells were immunostained with antibodies against tubulin. Images correspond to a maximum intensity projection of 6 slices, spaced by 1 micron. Scale bar correspond to 10 μm. Graph represent the total fluorescence intensity of tubulin per cell. Measurements were obtained from three independent experiments, for which n was between 10 and 20 cells for each cell type. Indicated P values were obtained by Mann - Whitney U-Test. C - Confocal images of MCF10A cells treated with siControl, siDSP, siEPPK, siPPK immnostained with antibodies against Vimentin to reveal the corresponding intermediate filaments. Images correspond to a maximum intensity projection of 6 slices, spaced by 1 micron. Scale bar correspond to 10 μm. Graph represent the total fluorescence intensity of vimentin per cell. Measurements were obtained from three independent experiments, for which n was between 10 and 20 cells for each cell type. Indicated P values were obtained by Mann-Whitney T-Test.

We then analyzed whether microtubule dynamics was associated to centrosome mispositioning in knocked-down cells since we have shown that microtubule stability participates to network polarization and centrosome off-centering toward inter-cellular junctions (Burute et al., 2017). Apart from centrosome mispositioning, we found no striking effect on the overall organization of the microtubules (Figure S3). We further tested microtubule stability by exposing cells to nocodazole for 5 min, and were surprised to found that the silencing of none of the three plakins had any effect on microtubule stability (Figure 5B).

Curious to identify which other parameter might impair microtubule network organization, we looked at intermediate filaments, which binds microtubules (Schaedel et al., 2021; Leduc and Etienne-Manneville, 2017). In particular vimentin filaments were shown to align with and template the spatial organization of microtubules (Gan et al., 2016). We thus stained for vimentin filaments and observed a massive increase of the amount of vimentin filaments in all plakin knocked-down cells (Figure 5C). Interestingly, the amount of vimentin is commonly used as a key EMT marker, suggesting that plakin knocked-down cells not only lost their epithelial polarity but might actually be engaged in a form of EMT.

## Conclusion

In this work we analyzed transcriptomic data with respect to two specific metrics: the polarity score and the EMT score of cell lines. We focused our analysis on the outcome of the proteins interacting with microtubules and found that several members of the plakin family stood out. Their expression levels were positively correlated to the level of epithelial polarity and negatively correlated with the EMT stage in two banks of mammary epithelial cells. We further tested experimentally the role of three plakins, epiplakin, periplakin and desmoplakin, by knocking down their expression with siRNAs in MCF10A. We confirmed that they were all involved in epithelial cell polarization.

Because of their size and diversity of cytoskeleton partners, plakins are called “giant cytolinkers” (Hu et al., 2018; Jefferson et al., 2004). They ensure the alignment of microtubules with intermediate filaments and bundles of actin filaments as well as their connections to cell adhesions (Bouameur et al., 2014; Suozzi et al., 2012; Prechova et al., 2023). They link them together and co-orchestrate their dynamics, notably during cell migration (Wu et al., 2008, 2011). Individually, at the interface with cell environment, or deeper within the cell cytoplasm, adhesions, actin filaments, intermediate filaments and microtubules are all essential players in the establishment of cell polarity (Ebnet et al., 2018; Oriolo et al., 2007; Li and Gundersen, 2008). Our work suggests that, by transmitting structural information from peripheral adhesions to inner cytoskeleton networks, plakins ensure a functional coherence between cell environment and cell interior in the organization of epithelial cell polarity.

Plakins not only connect cytoskeleton filaments to plasma membrane, they also bind them to the nuclear membrane (Liem, 2016). This connection might have a structural role in the organization of the cytoplasm, but it may also be part of a regulatory pathway, since mechanical forces on the nucleus affect gene expression (Uhler and Shivashankar, 2017). Indeed, plakins contribute to reorganize cytoskeleton networks during embryo development and cell differentiation (Sonnenberg and Liem, 2007). In this study, we revealed their role in cell polarity by searching for proteins participating to the destabilization of epithelial polarity during EMT. Their expression appeared anti-correlated to EMT, but they might also have a causal role in this transition. Indeed, in all the three investigated plakins knocked-down cells, we observed an overexpression of vimentin, which is a well-established marker of the mesenchymal phenotype. So more than destabilizing the epithelial polarity, their disappearance might instruct the acquisition of the mesenchymal phenotype. This would be consistent with their frequent mutation (Hu et al., 2018), and their decreased expression (Wesley et al., 2021) in various cancers. Thus, these giant cytolinkers that co-organize all cytoskeleton components, might also act as signaling platforms that instruct gene expression and direct cell differentiation. This possible regulatory role of plakins in cell differentiation raises the hypothesis that cytoskeletal organization is linked to protein expression profiles to ensure functional coherence between cell architecture and identity.

## Acknowledgements

The authors thanks Emilie Henry and Cecile Reyes from the Genomics platform of Institut Curie for STR analysis and microarray experiments, and Dr Pierre de la Grange from GenoSplice for gene expression analysis of microarray data. We thank Elaine Del Nery and Aurianne Lescure for the siRNA screen we did together on the BioPhenics platform, which contributed to the identification of plakins although these data could not be included in this manuscript. This work was supported by the European Research Council: Consolidator Grant 771599 and Proof-of-Concept Grant 780458 to Manuel Théry and an Advanced Grant 741773 (AAA) to Laurent Blanchoin.

## Authors contribution

Juliana Geay performed all experiments relative to the validation of plakins in MCF10A (Figure 4 and 5). Yoran Margaron performed the experiments relative to the polarity of TNBCs (Figure 1). Laurent Guyon analyzed the transcriptomic data and computed the EMT score (Figure 1, 2 and 3). David Gentien and Fabien Reyal provided the TNBC and their transcriptomic data. Alain Puisieux provided the transcriptomic data of HME. Manuel Théry and Laurent Blanchoin obtained the fundings and co-supervised the study. Manuel Théry wrote the manuscript.

## Material and Method

### Transcriptomic analysis

TNBC cell lines freshly obtained from new vials of the ATCC supplier were expanded following ATCC recommendations. Large cell preparation were first used for DNA preparation and Short Tandem Repeat (STR) analysis to confirm cell authentication. Transcriptomic analysis were conducted on total RNA purified with miRNeasy kit. A quality control of total RNA was carried out with a Nanodrop ND1000 spectrophotometer (Thermo Fisher) to monitor the concentration and purity of samples, and integrity of total RNA was controlled using RNA6000 Lab-on-a-chip with a Bioanalyzer (Agilent technologies). Transcriptomic analysis was based on Affymetrix Human Exon 1.0 ST Array hybridization following the supplier recommendations, as published before (Lerebours F, et al. 2020). A data quality control was realized with Affymetrix Expression Console. Gene expression analysis was carried out with EASANA (GenoSplice Technology), and GenoSplice’s FAST DB annotations.

Data were normalized using the RMA normalization procedure to create quantile - normalized log2 transcript signal values. The transcriptomic data for the HME cell lines were already published in (ref (Morel et al., 2012) à rappeler ici), and made available by the authors (GSE32727). The expression of different probes for a given gene were averaged to end up with a single expression per gene.

Transcriptomic data and correlations were analyzed using R version 4.2.1 (R Core Team, 2013).

### EMT Score

Taube and collaborators provide a signature of genes overexpressed or repressed following EMT (Taube et al., 2010). In this list, the 239 genes expressed in the HME cells correspond to 150 repressed genes following EMT in the Taube *et al*. list and 89 overexpressed genes. An EMT score was computed per cell line by averaging the expression of the 89 genes minus the average of the expression of the remaining 150 genes. The higher the score, the more advanced the transition from epithelial to mesenchymal state.

### Gene Ontology enrichment

To extract biological meaning from the list of genes for which the expression is correlated to either polarity or EMT, we performed gene ontology enrichment with DAVID (Huang et al., 2009). We selected the genes for which the correlation is above +0.5 or below −0.5 to perform the enrichment. We only provide the significant ontologies with a threshold of 0.05 after correction for multiple testing with the Benjamini and Hochberg procedure.

### Cell culture

BT20, MDA-MB-157, MDA-MB-231, MDA-MB-436 and MDA-MB-468 cell lines were cultured in Dulbecco’s Modified Eagle Medium (31966, Gibco) supplemented with 10% FBS (50900, Biowest) and 1% antibiotic-antimycotic (15240-062, Gibco). BT-549, HCC38, HCC70, HCC1143, HCC1937 were cultured in RPMI 1640 (61870, Gibco) with 10% FBS and 1% antibiotic-antimycotic. MCF10A cells comes from ATCC bank of cells. Cells were grown in MEBM medium (Lonza) supplemented with MEGM kit (Lonza) and 100 ng/mL choleratoxin (Merck) at 37°C and 5% CO2.

### Micropatterning

Coverslips were cleaned with acetone and isopropanol before plasma-cleaned for 5 min and incubated with 0.1 mg/ml poly-lysine/polyethylene glycol (PLL-PEG) diluted in 10 mM HEPES for 30 min at room temperature. Excess PLL-PEG was washed off coverslips using distilled water, and the coverslips were then dried and stored at 4 °C overnight before printing. Micropatterns were obtained on prepared PLL-PEG coverslips by exposing them to deep UV for 5 min through a designed chrome masks as described previously (Azioune et al., 2010) and coated with 50:50 (v/v) fibronectin/fluorescent fibrinogen mixture (20 µg/ml each) and collagen 5 µg/ml diluted in fresh 100 mM NaHCO_3_ (pH 8.3) for 30 min at room temperature. Micropatterned coverslips were washed twice in NaHCO_3_, and incubated for 30 min with culture medium before use. Plated cells (approximately 30000 cells/ml cells) were allowed to adhere for 24 h before fixation. H shaped patterns have an area of 1100 µm².

### Immunofluorescence

Migrating cells (for 8 h), sparse or a monolayer of cells, were either fixed with 4% PFA, 0.25% Triton, 0.2% Glutaraldehyde 25uM in cytoskeleton buffer for 10 min at 37°C, washed thrice with PBS, and quenched with Sodium Borohydride for 10 min at 4°C, or fixed with ice-cold methanol for 3 min at −20°C. Cells were washed thrice in PBS and stored at 4°C. Coverslips were blocked for 30 min with 5% BSA in PBS. Primary and secondary antibody were incubated 1 h at room temperature in 5% BSA. Coverslips were incubated 10 min with DAPI. Coverslips were mounted with Mowiol mounting medium (81381, Sigma). Fluorescence images were acquired with a Spinning Disk microscope equipped with 40× 1.25 NA or 63× 1.4 NA objectives and recorded on a SCMOS camera (BSI) with Metamorph software

### Antibodies

Primary Antibodies used in this study are: Pericentrin (1 : 1000, ab4448, rabbit polyclonal, Abcam), anti-α-tubulin (1 : 400, ab18251, rabbit polyclonal, Abcam), Vimentin (D21H3, 1 : 400, 5741S, rabbit monoclonal, Cell Signaling Technology), Alexa Fluor 555 Phalloidin (1 : 400, A34055, Invitrogen), Alexa Fluor 647 Phalloidin (1 : 400, A22287, Invitrogen), anti - Desmoplakin (1:500, ab16434, mouse monoblonal, Abcam), anti-Epiplakin (1:500, ab247172, polyclonal rabbit, Abcam) and anti-Periplakin (1:500, ab72422, rabbit polyclonal, Abcam).

Secondary antibodies used were: Alexa Fluor 488 Goat anti-Rabbit (A-11008), Alexa Fluor 568 Goat anti-Rabbit (A-11036), Alexa Fluor 488 Donkey anti-Mouse (A-21202), Alexa Fluor 555 Donkey anti-Mouse (A-31570), Alexa Fluor 647 Donkey anti-Mouse (A-31571) from Invitrogen.

### siRNA

MCF10A cells were transfected using Lipofectamine RNAiMAX (13778150, ThermoFisher Scientific) according to the manufacturers’ protocol. For siRNA gene silencing siRNA sequences used were:

Luciferase (control) 5’-UAAGGCUAUGAAGAGAUAC-3’,

Desmoplakin 1 5’-CCGACATGAATCAGTAAGTAA-3’,

Desmoplakin 2 5’-CAGGGAGATCATGTGGATCAA-3’,

Epiplakin 1 5’ - CCGGCTGACCGCCATCATCGA-3’,

Epiplakin 2 5’ - CACGCAAGAGAAGGTCTCGTA-3’,

Periplakin 1 5’ - CAACCGGAACCTGGAGGCCAA-3’,

Periplakin 2 5’ - CTGAGGCCCGTGAGAAGGTAA-3’.

siRNAs were used at 1 nmol and experiments were carried out 3 days after transfection.

### Western Blot

Cell lysates were obtained with SDS supplemented with Laemmli Buffer 2x (Sigma). Samples were boiled for 10 min at 94°C before loading on polyacrylamide gels (4-15% Bis-Tris Gel, Bio-Rad). Proteins were separated by gel electrophoreses at 80 V and transferred at 90 V for 2 h on nitrocellulose membranes (Precast Protein Gels,ref 4561084 and 0.2um nitrocellulose membrane, ref 1620112). Membranes were blocked with a solution of 5% BSA in Tris-buffered saline, 0.1% Tween 20 detergent (TBS-T). The membranes were incubated for 1 h at room temperature with primary antibody, and overnight at 4°C with horseradish peroxidase (HRP) - conjugated secondary antibody. Both primary and secondary antibody solutions were made in a solution of 5% BSA in TBS-T. Protein bands were revealed with ECL chemoluminescent substrate (Biorad) and signals were recorded using a ChemiDoc MP Imaging System (Biorad). The following primary antibodies were used for western blot analysis: anti-GAPDH (1:10000, sc-25778, rabbit polyclonal, Santa Cruz), anti-Desmoplakin (1:500, ab16434, mouse monoclonal, Abcam), anti-Epiplakin (1:500, ab247172, polyclonal rabbit, Abcam) and anti - Periplakin (1:500, ab72422, rabbit polyclonal, Abcam). Secondary antibodies used were HRP donkey anti-rabbit and goat anti-mouse (both 1:1000, Invitrogen).

### Images analysis

#### Polarity Index

Image processing was performed using an ImageJ macro. First, fluorescent cell-adhesive patterns were individualized using a template matching method. Then nucleus and centrosome detection were realized based on threshold and size filtering within each cell of the doublet. Taking into account that only cell doublets were considered for further analysis, groups of cells encompassing two nuclei and two centrosomes were manually selected. The center of mass of the nucleus was computed and each centrosome was assigned to the closest nucleus. We then computed the coordinates (x,y) of the centrosome with respect to the nucleus.

#### Polarity score

The percentage of highly polarized cells was defined as the proportion of cells having a centrosome located in close proximity of the cell-cell junction, in a triangular area defined as: |*y*| ≤ (*x* − 0.5) ∗ tan 60, with 0.5 ≤ *x* ≤ 2.

Vimentin intensity cell per cell was measured using Fiji by detecting cell contour and using the plugin “measure” to obtain the total fluorescence intensity.

## Statistical analysis

Statistical analyses were performed using GraphPad Prism software (Version 5.00, GraphPad Software). For each experiment, cell sampling and the number of independent replicates is indicated in figure legends. Data sets with normal distributions were compared with one-way Anova test. Non normal distributions were compared with a Mann-Whitney test.

---

**Figure S1.**
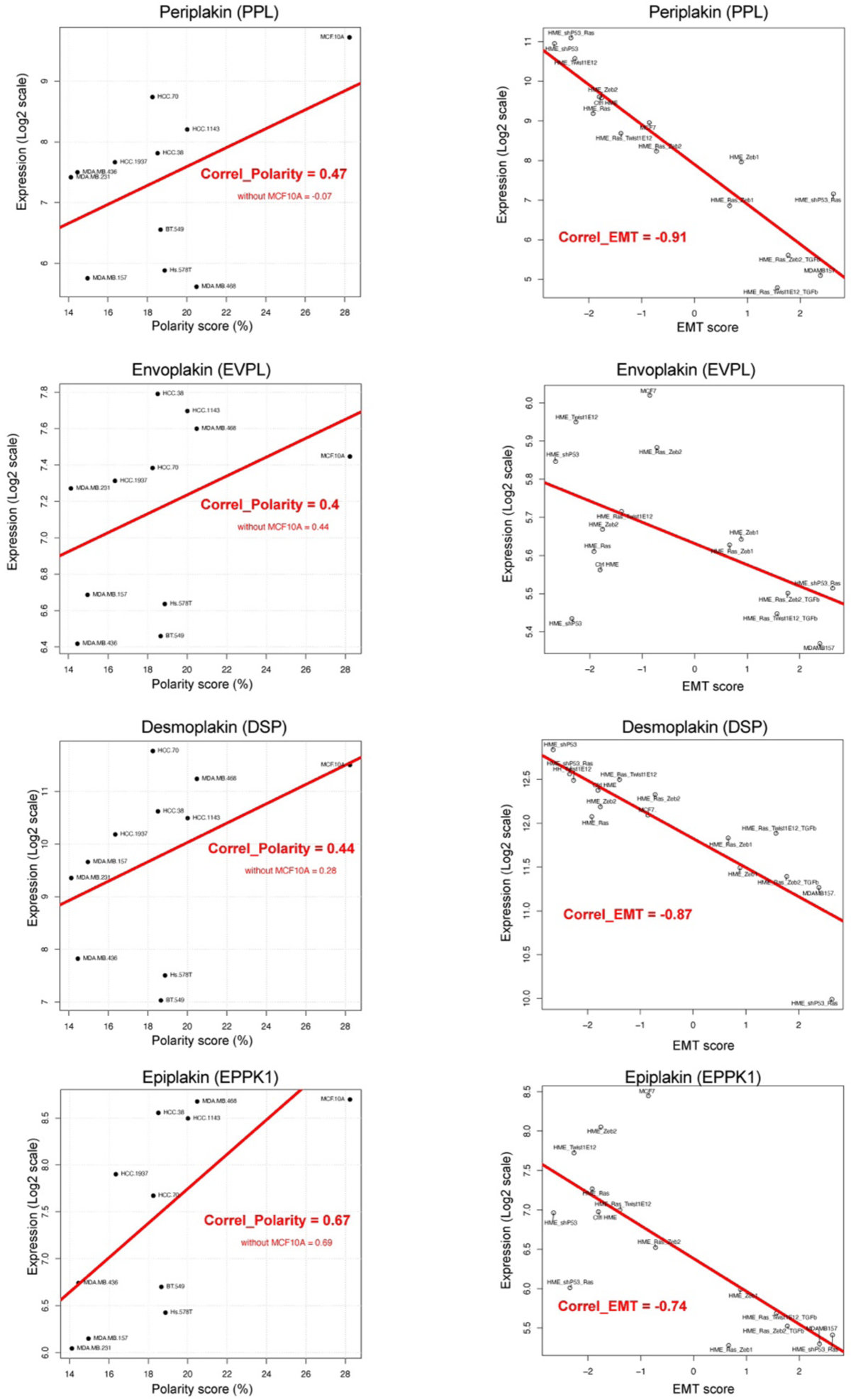
Correlation of the expression levels on plakins with the Polarity and the EMT score These data are associated to figure 1 and 2. They show the relationship between the level of expression of plakins and the polarity score of TNBC (left) or the EMT score of HME cells lines (right).

**Figure S2.**
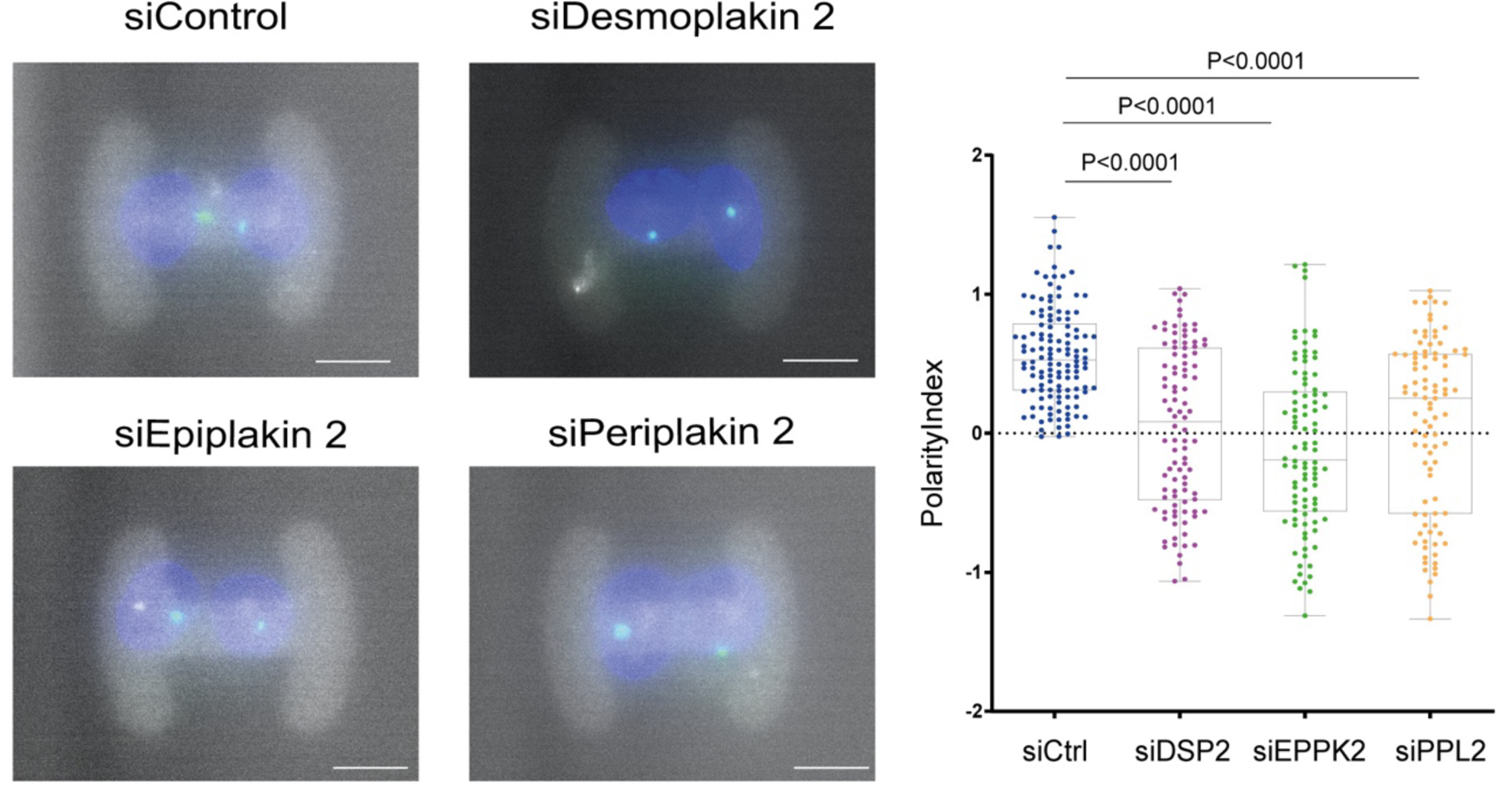
Plakins and cell polarity These data are associated to Figure 4. They show the same type of measurement of cell polarity as those shown in Figure 4D but cells were treated with the second siRNA identified shown in Figure 4A.

**Figure S3.**
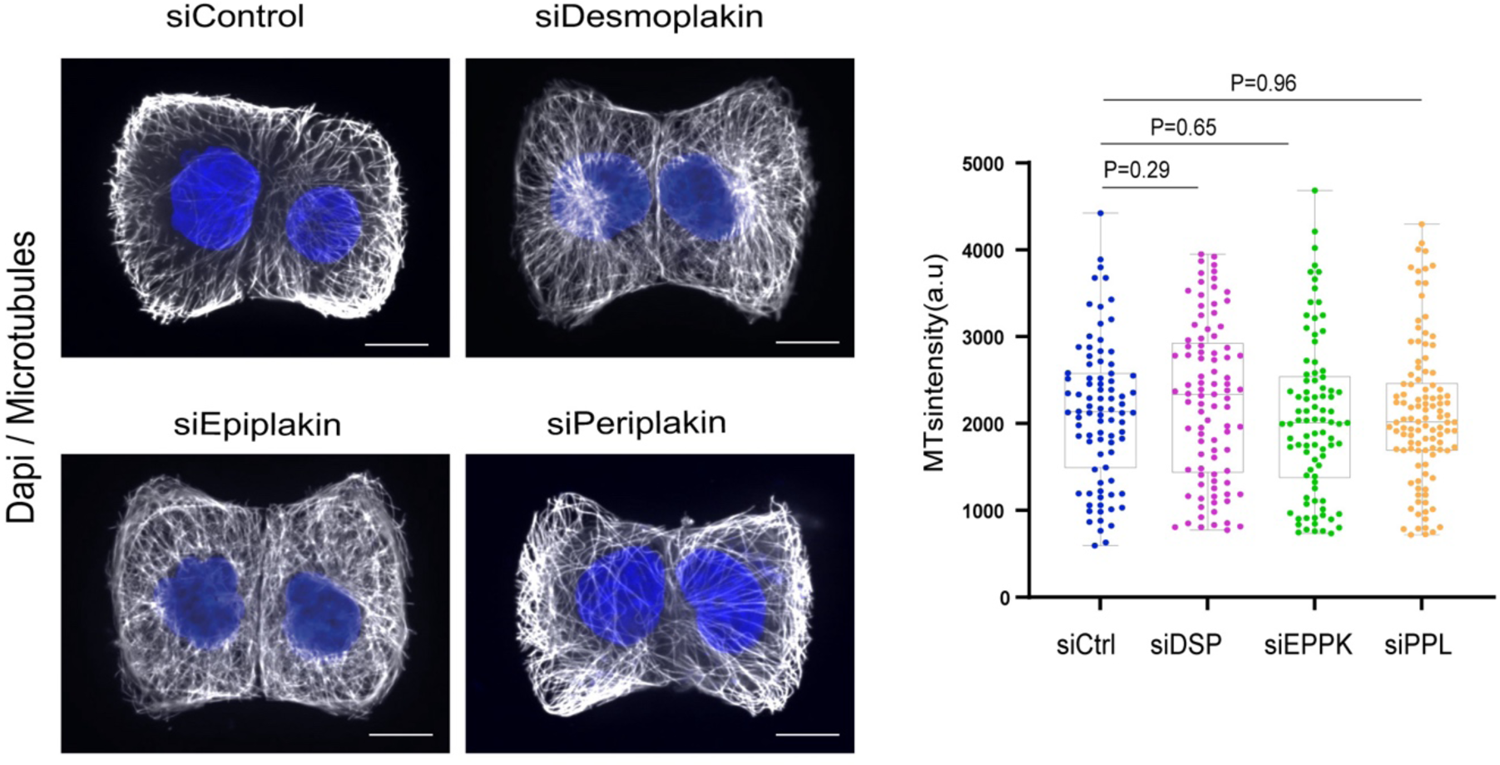
Plakins and microtubules These data are associated to figure 5B. They show confocal images of MCF10A cells treated with siControl, siDSP, siEPPK, siPPK. Cells were fixed and immunostained with antibodies against tubulin. Images correspond to a maximum intensity projection of 6 slices, spaced by 1 micron. Scale bar correspond to 10 μm. Graph represent the total fluorescence intensity of tubulin per cell. Measurements were obtained from three independent experiments, for which n was between 10 and 20 cells for each cell type. Indicated P values were obtained by Mann - Whitney T-Test.

